# Control of filament length by a depolymerizing gradient

**DOI:** 10.1101/2020.06.02.130039

**Authors:** Arnab Datta, David Harbage, Jane Kondev

## Abstract

Cells assemble microns-long filamentous structures from protein monomers that are nanometers in size. These structures are often highly dynamic, yet in order for them to function properly, cells maintain precise filament lengths. In this paper, we investigate length dependent depolymerization as a mechanism of length control. This mechanism has been recently proposed for flagellar length control in the single cell organisms Chlamydomonas and Giardia. Length dependent depolymerization can arise from a concentration gradient of a depolymerizing protein, such as kinesin-13 in Giardia, along the length of the flagellum. Two possible scenarios are considered, that of a linear and an exponential gradient of depolymerizing proteins. We compute analytically the probability distributions of filament lengths for both scenarios and show how these distributions are controlled by key biochemical parameters through a dimensionless number that we identify. In Chlamydomonas cells it has been well documented that the assembly dynamics of its two flagella are coupled via a shared pool of molecular components, and so we investigate the effect of a limiting monomer pool on the length distributions. Finally, we compare our calculations to experiments. While the computed mean lengths are consistent with observations, the noise is two orders of magnitude smaller than the observed length fluctuations.

## I. Introduction

All eukaryotic cells contain nanometer-sized proteins that polymerize to form filaments that are hundreds of nanometers to tens of microns in length. These filaments often associate with other proteins to form larger structures that perform critical roles in cell division, cell motility, and intracellular transport. They are often highly dynamic, experiencing a high turnover rate of their constitutive protein subunits[1,2]. Despite this, in order to function properly some of these structures, like the mitotic spindle, or actin cables in budding yeast[2,3] must maintain specific and well-defined sizes.

When considering dynamics of filament length, the two basic processes are the addition of monomer units, characterized by the rate of assembly, and the removal of these units, which is described by the rate of disassembly. For the dynamics to reach a steady state with a well-defined length, one or both rates must be length dependent. Therefore, the search for the molecular mechanism of length regulation often boils down to finding out whether the rate of assembly or disassembly (or both) is length dependent, and what molecular-scale interactions produce this length dependence[4,5].

Many mechanisms for length regulation have been proposed based on careful experiments in cells and on reconstituted filamentous structures in vitro[3,6–9]. One well studied mechanism [10] of length control is to limit the number of subunits available for assembly. For example, in vitro experiments have demonstrated how the size of the mitotic spindle depends on the size of the compartment in which it self-assembles, consistent with a limiting pool mechanism for length control[11]. In the case of multiple structures all assembled from the same pool of components this mechanism can only control the total number of subunits in all the structures, but is incapable of controlling the size of each individual structure[12]. How cells create and maintain multiple filamentous structures, such as cilia in a multiciliate cell, is the key question that motivates this paper.

The question of length control of multiple filaments that share a monomer pool has been studied extensively in the single cell algae Chlamydomonas reinhardtii (“Chlamydomonas”)[13–15]. There, experiments have directly demonstrated that the assembly dynamics of the two flagella are coupled. If one flagellum is cut, thereafter the remaining flagellum shrinks while the cut flagellum grows, until the two reach the same length, which is smaller than the length before the cut. Once the lengths of the two flagella equalize, they continue growing at the same rate until the original lengths are reached, and the lengths are stable. In experiments where drugs are added that prevent protein production, after both flagella are cut, they regrow to a length that is shorter than the length before the cut. These experiments indicate that the free tubulin pool is limiting, or some other protein required for assembling a flagellum is the limiting component, and is shared between the two flagella [15]. Furthermore, these experiments imply the presence of a length control mechanism that is able to detect the lengths of individual flagella, which the limiting pool mechanism cannot do as it is only sensitive to the total number of monomers in both flagella [12].

The mechanism of length control of the two Chlamy flagella that has thus far received the most attention is one that focuses on length dependent assembly [16,17]. Recently we have shown that length dependent assembly can arise from a limited and shared pool of motors that transport tubulin to the flagellar tip, where assembly occurs [18]. In this case, though, the assembly rate for the two flagella is the same and depends on the sum of their lengths. We showed that this mechanism still fails to control individual filament lengths and proposed instead that length dependent depolymerization might be the key length controlling process. Moreover, in a different single cell organism, Giardia lamblia, which has four pairs of flagella of different length, recent experiments have shown that the assembly rate is independent of flagellar length, also suggesting length-dependent depolymerization as the mechanism of length control [19].

Examples of length-dependent depolymerization are abundant in the literature. In vitro studies of kinesin-8 have directly demonstrated that its rate of depolymerization scales linearly with the length of the microtubules it binds to[20]. Other, in vivo studies have demonstrated that the microtubule depolymerizing kinesin-13 localizes to the tips of flagella, and that reduction of its expression in the cell leads to longer flagella [21–23]. In Giardia this localization was shown to be quantitatively consistent with directed transport of kinesin-13 to the tip of the flagellum, that in combination with the depolymerizing activity of this non-motile kinesin leads to a length dependent depolymerization[19].

In this paper we examine how a gradient of a depolymerizing protein along the length of a filament, can control the length of multiple filaments via length dependent depolymerization. Using a master equation for the lengths, we show that length dependent depolymerization can produce well defined steady state lengths for multiple filaments, even when the filaments lengths are coupled by a limiting pool of monomers. The resulting length distributions are coupled, and we calculate the joint probability distribution by solving the master equation in the steady state. For parameter values typical for a Chlamydomonas cell, our model produces steady state filament lengths comparable to the flagella in these cells.

## II. Depolymerization gradient

Length dependent depolymerization has been shown to occur when a depolymerizing factor is actively transported to the ends of filaments by the action of motors [3,19,20]. The length dependence of the depolymerization rate arises due to the formation of a spatial gradient of the depolymerizing factor along the length of the filament (see figure 1). Quantitative measurements using fluorescent labeled proteins that are transported along a filament have uncovered gradients that vary linearly [20] and exponentially [19,24] with the length of the filament. Here we consider both types of concentration gradients of depolymerizing factors as a mechanism of length control in flagella.

**Figure 1.**
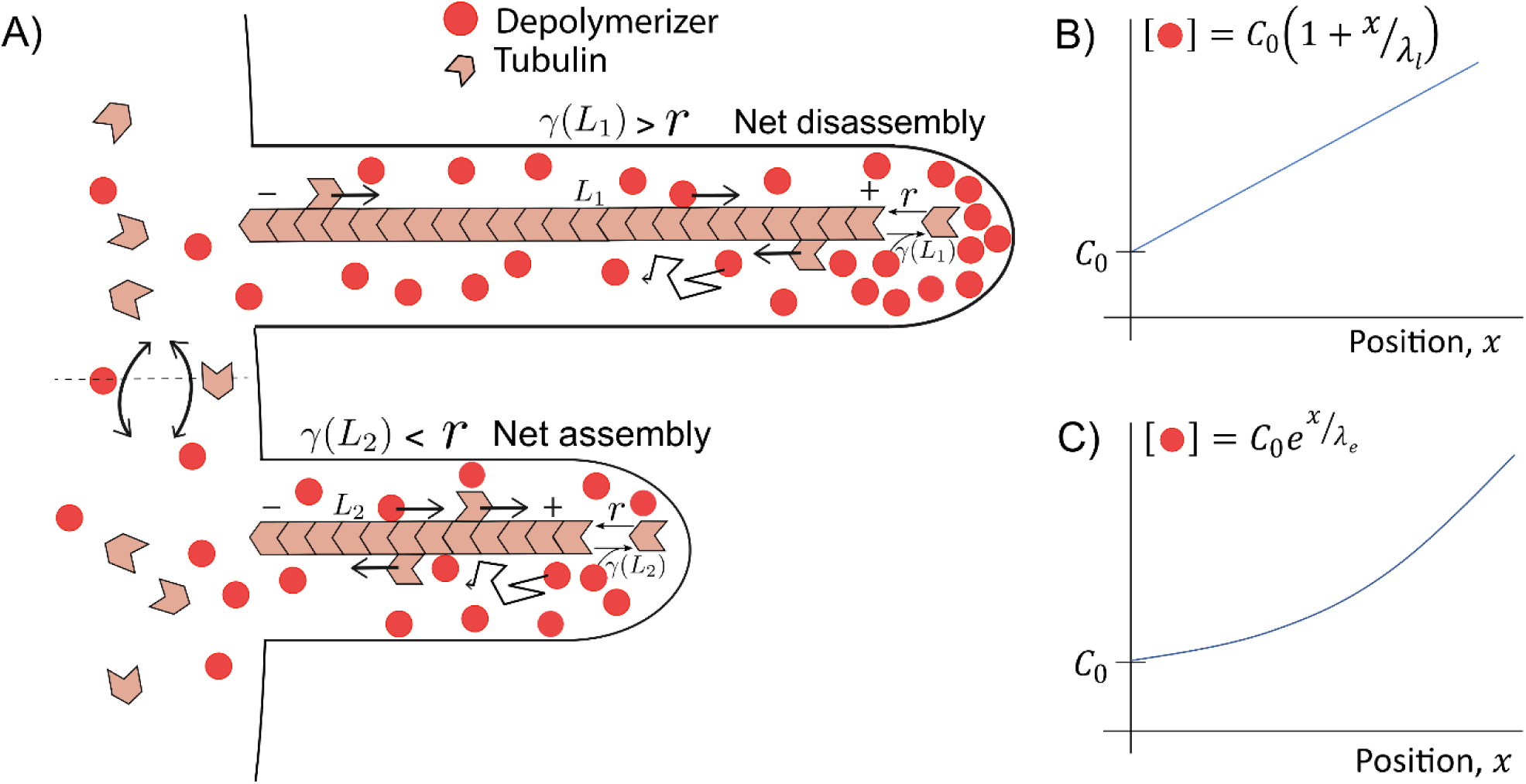
Schematic of gradient formation in flagella. **A.** Depolymerizing proteins (red) gather at the end of the filament, either by directed transport or by diffusion. The resulting length dependent gradient determines the rate of disassembly, after which the free IFT particles and the monomers diffuse back to the cell body. The assembly rate is determined by the monomer pool in the cell body. **B.** Shows the linear concentration gradient, which forms when the IFT particles can only bind to the filament base. **C.** Shows the exponential concentration gradient which forms when the IFT particles can bind anywhere on the filament. In both **B** and **C** the bound IFT particles can detach only at the filament tip.

Linear and exponential gradients arise from a simple mechanism of combined directed transport and diffusion of molecules within a flagellum, as shown in figure 1. In both cases the depolymerizer is transported by motors toward the tip of the flagellum, where it is released. In case when the loading of the depolymerizer only occurs at the proximal end of the flagellum, 

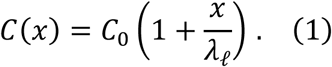

Here, *C*_0_ is the concentration of depolymerizers in the shared pool and *λ*_*ℓ*_ is the length scale of the gradient. Such a gradient was observed directly in experiments with fluorescently labeled kinesin-2 in Chlamy, which are loaded at the proximal end of the flagellum, move directionally toward the distal end, and return back to the base diffusively [25]. These motor proteins are responsible for the anterograde motion of the IFT particles, and whether other IFT proteins show a similar concentration gradient, is not known.

In the second case we consider, the depolymerizer is free to bind to the motors that are moving along the filament anywhere in the flagellum. (Note that whether the depolymerizer binds to a motor directly, or to cargo carried by the motor, does not affect the mathematics of the model.) The concentration gradient of the depolymerizers, in this case is given by, 

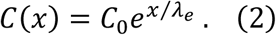

Once again *C*_0_ is the concentration of depolymerizer of depolymerizers in the shared pool and 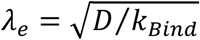 is the length scale of the gradient. *D* is the diffusion coefficient of the depolymerizer in the flagellum and *k*_*Bind*_ is the rate with which it binds to the motors moving along the filament[19,24].

The presence of a concentration gradient of a depolymerizing protein leads to length dependent depolymerization. Namely, we assume that disassembly of the filaments only happens at the tip of the filament, *x* = *L*, as is observed in Chlamy[26–29]. For the depolymerizer to remove a monomer from the tip of the filament, it must first diffusively find the end of the filament, bind to it, and then remove a subunit. We assume that depolymerizer concentration is low, so that finding the end of the filament is the rate limiting step. In that case the rate of depolymerization is proportional to the concentration of the depolymerizer at the tip, 

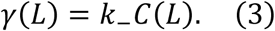

For example, in vitro studies of MCAK, a depolymerizing non-motile kinesin, show this kind of linear relation for MCAK concentrations less than 10 nM [22].

We consider the polymerization rate to be proportional to the concentration of free monomers in the shared cytoplasmic pool from which they are drawn into the flagellum by directed transport of the motors (see figure 1A), 

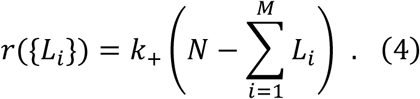

Here *N* is the total number of monomers, *M* is the number of filaments, and *L*_*i*_ is the length of the *i*^th^ filament in monomer units (for example, one micron of a microtubule corresponds to roughly 2000 monomers, which are, in this case, tubulin dimers [30]). We assume that the assembly rate per monomer (*k*_+_) is length independent, so all the length dependence of assembly comes from the pool of monomers being limited. In particular, we do not take into account the length dependence of *k*_+_ that can arise due to a limited pool of motors that transport monomers to the tip of the flagellum (Figure 1), which was analyzed in ref.[18]. There it was shown while this feature of the assembly dynamics is important for getting the regeneration dynamics to agree with experiments in Chlamy, but it does not provide length control for multiple flagella. Giardia experiments [19] on the other hand found no such length dependence of *k*_+_, which is another reason we choose to leave it out of our analysis in this paper.

In Chlamy, experiments suggest that a substantial fraction of the tubulin in the cell is in the two flagella, and the presence of other microtubule structures within the cell body further decrease the amount of free tubulin [15,31,32]. In Giardia the tubulin pool seems to be much larger as there is no evidence for its depletion [19], therefore, the polymerization rate is likely length-independent. Below, we consider both cases and show that length dependent depolymerization leads to filaments with a well-defined length, regardless of whether the polymerization rate is length dependent or not.

The model we have adopted for the dynamics of flagella describes each flagellum as a single filament and does not take into consideration the fact that the flagellum is composed of from multiple microtubule doublets and other structural components [16]; we take this structure into account only in the sense that each monomer that attaches or detaches to the end of the filaments contributes to a change in length that is only a small fraction of the monomer size (as estimated below, we estimate that a micron of the flagellum takes up 20,000 tubulin dimers).

## III. Length control by length dependent depolymerization

We use the length-dependent disassembly model described above to compute the steady state filament lengths and the fluctuations around the mean in steady state. The steady state length is obtained by finding the filament length for which the rate of polymerization and depolymerization are equal. In order to characterize the length fluctuations in steady state, we make use of the master equation for the probability distribution of filament lengths.

### A. Large monomer pool

First, we consider the case when the monomer pool is large enough so that in steady state the monomers taken up by the filaments are only a small fraction of the total number of monomers (*N*) in the cell, Σ_*i*_*L*_*i*_ ≪ *N*. In this case we can approximate the assembly rate as a constant, independent of length *r* ≈ *k*_+_*N*; *k*_+_ is the rate of assembly per monomer. Recent experiments in Giardia found a length independent polymerization rate for its flagella, consistent with this assumption. Furthermore, in this case the dynamics of individual filaments to be uncoupled and we can restrict our analysis to an individual filament.

In steady state the rate of polymerization and depolymerization are equal. Using equations (3) and *r* ≈ *k*_+_*N* for the assembly rate, we find the steady state length 

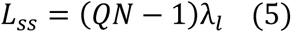

where we have used equation (1) for the concentration of the depolymerizer at the tip, which corresponds to the case of a linear depolymerization gradient. Here we have introduced a dimensionless quantity 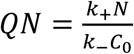. This quantity is the ratio of the rate of polymerization and the rate of depolymerization, when the concentration of depolymerizing proteins is equal to the cytoplasmic concentration *C*_0_. As will become clear later, this is the key dimensionless quantity that controls both the steady state filament length and the length fluctuations.

Similarly, for the case of the exponential gradient of depolymerizers, using equation (2), we find 

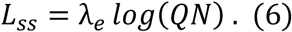

From equation (5) and (6) we see a clear difference in the way in which the linear and exponential gradients of depolymerizers determine the steady state filament length. In the case of the exponential gradient, equation (6), the filament length is set by the length scale of the gradient *λ*_*e*_; for reasonable values of the parameters (discussed below), *log*(*QN*) ≈ 1. This is unlike the case of the linear gradient, where the filament length is proportional to *QN*, and therefore the steady state length is much more sensitive to the chemical rate constants, as well as the overall concentration of depolymerizers and monomers in the cell.

To quantify the precision of a length control mechanism we compute the fluctuations in length around the steady state value using the master equation 

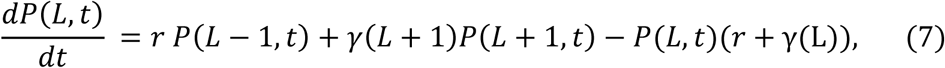

where *P*(*L, t*) is the probability that at time *t* a filament has length *L*. Once again, the length dependent depolymerization rate *γ*(*L*) is given by equation (3), while the polymerization rate is constant and given by *r* = *k*_+_*N*.

For the linear depolymerizer gradient we find the steady state probability distribution, 

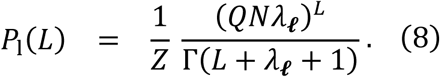

where 1/*Z* is the normalization constant and Γ is the gamma function. The approximate mean length of the distribution is, ⟨L⟩_l_ ≈ (*QN* − 1)*λ*_***ℓ***_ which is also the steady state length; see equation (5). The variance of the distribution is *var*_*l*_*L* ≈ *QNλ*_***ℓ***_. Both the mean and the variance are dimensionless as *λ*_*l*_ is given in monomer units. (We have also derived exact expressions for the mean and the variance, but they are practically identical to the approximate ones for *QN, λ*_***ℓ***_ ≳ 2, which is always the case.)

For the exponential depolymerizer gradient we find that 

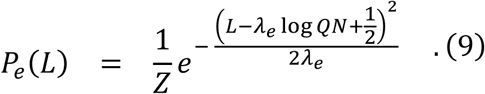

where to a good approximation the normalization constant is 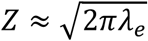. In this case, the approximate mean length is, ⟨*L*⟩_*e*_ ≈ *λ*_*e*_ Log *QN* and the variance *var*_*e*_*L* = *λ*_*e*_; as above, the length scale of the gradient, *λ*_*e*_, is expressed in units of monomers.

To make numerical estimates of the parameters that define the length distributions we make use of published data on flagella in Chlamy cells, whose steady state length is *L*_*ss*_ ≈ 10*μm*. Since each micron of a microtubule contains ≈ 2000 tubulin dimers and each flagellum consist of roughly 10 microtubules, we estimate that there are around 2 × 10^4^ tubulin dimers per micron of a flagellum. Therefore, a typical flagellum of length 10 microns contains 2 × 10^5^ tubulin dimers. It has been estimated [31] that around 20% of the total amount of tubulin in the cell is used to make the flagella, so we estimate that the total number of tubulin dimers in the cell, *N* = 2 × 10^6^.

In ref.[25] a linear gradient of kinesin-2 proteins was measured along the length of the Chlamy flagella and this data provides an estimate of *λ*_***ℓ***_ ≈ 3 μ*m*, for the characteristic length of the gradient. According to our finite pool model (discussed below), for the linear gradient model and assuming a steady state length of 10 microns, gives QN ≈ 5.

The dimensionless parameter 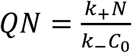 is a ratio of the maximum polymerization rate (when all *N* monomers are cytoplasmic) and the depolymerization rate at the cytoplasmic concentration of the depolymerizing molecule (*C*_0_). Based on experiments in which regrowth of a Chlamy flagellum is monitored after having been cut, we estimate the maximum polymerization rate to be a few microns a minute. Measurements of the depolymerization rate of MCAK [22] find that depolymerization of a few microns a minute is achieved at nanomolar concentration. Assuming that this is a good estimate for the depolymerization in the cell, we find *QN* ≈ 1 for depolymerizer concentration (*C*_0_) of a few nanomolar. If the actual cytoplasmic concentration of depolymerizer is in the picomolar range (which would correspond to tens of molecules of the depolymerizer in the Chlamy cell, which is tens of microns in diameter), we would end up with the estimate *QN* ≈ 1000. Therefore, we take 1 < *QN* < 1000 as a reasonable range of values for this dimensionless number. For example, in figure 2, we have chosen *QN* = 200, which for the exponential gradient model with a finite pool of monomers (discussed below) and a steady state flagellum length of ten microns, gives an estimate *λ*_*e*_ ≈ 4 × 10^4^ monomers or approximately 2 μ*m*.

**Figure 2:**
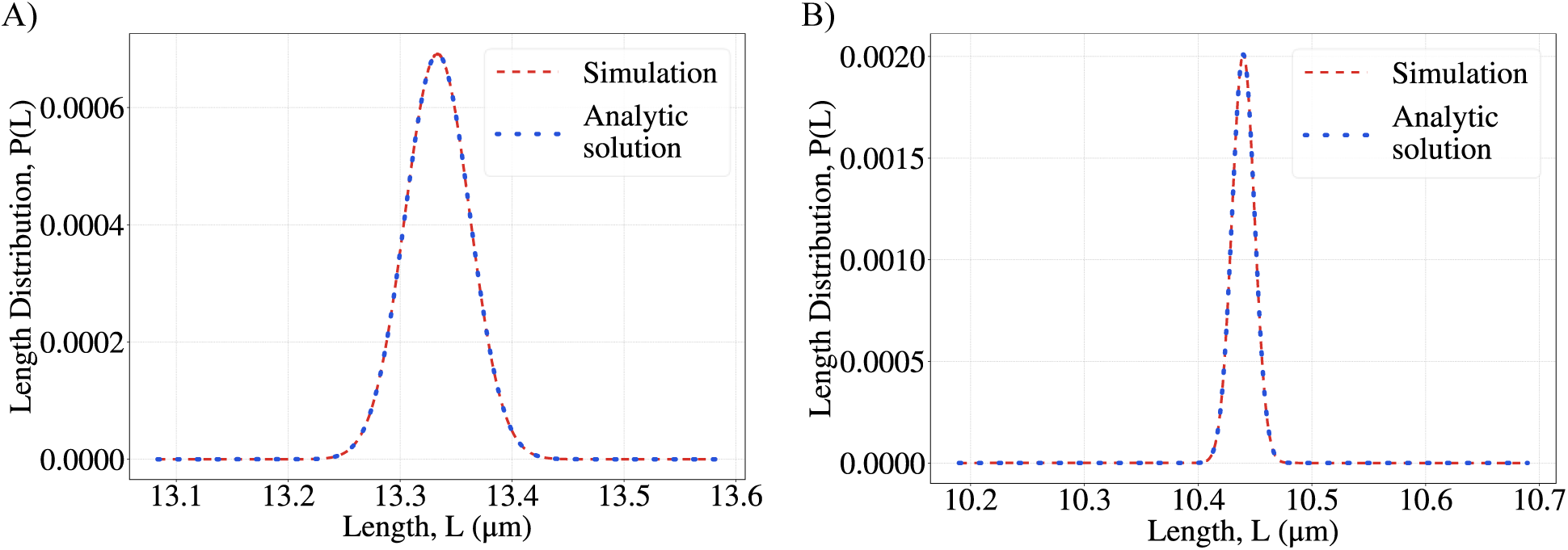
Probability distributions of the filament lengths for **A.** the linear depolymerizer gradient model and **B.** the exponential gradient model. Parameters: *N* = 2 × 10^6^. For **A**. *QN* = 5, *λ*_*l*_ = 6.66 × 10^4^(in monomer units). For **B.** *QN* = 200, *λ*_*e*_ = 3.94 × 10^4^ (in monomer units).

In order to characterize the length fluctuations, we also compute the noise for each model, 

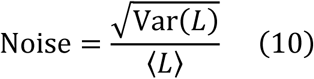

and its dependence on the model parameters. For the linear gradient model, using equation 8 we find 

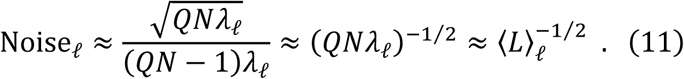

Therefore, as the average length increases, the noise decreases and the length of the filament becomes better defined.

Using equation (9) for the length distribution in the exponential gradient model, we find 

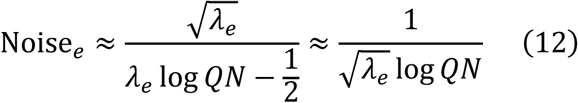

and, once again, the model predicts that the length will become better defined as it increases.

If we compare our results for the noise in figure 2, to the experimentally measured length fluctuations[15] we see that the theoretically determined noise is ≈ 100 times less than the observed noise. Therefore, the observed length fluctuations cannot be solely due to the stochasticity of the assembly dynamics described by our models.

### B. Limiting pool of monomers

The two models investigated in the previous subsection both describe the stochastic assembly of filaments that produce a well-defined length, and unlike the limited subunit-pool model [12,18] they can simultaneously regulate the lengths of any number of filaments. This is because the dynamics of filament assembly are independent of one another. Here we examine the situation when the length dynamics of multiple filaments are coupled via a shared and limiting pool of subunits.

For example, experiments on Chlamydomonas cells have shown that the assembly of its two flagella are not independent [15]. As described in the Introduction, when one flagellum is cut, the length of the remaining flagellum decreases in response, while a new flagellum grows in place of the one that was removed. This observation supports the idea that the pool of tubulin used to assemble the flagella, is limited. Furthermore, it has been determined that in Chlamydomonas more than half of the tubulin in the cell can be in its two flagella [15,31,32], which also suggests that the tubulin pool is limited.

The key question we address now is, how does the coupling of the filament-length dynamics via a shared and limited pool of monomers affect the mean and the variance of the length of each individual flagellum? We consider the master equation for two filaments: 

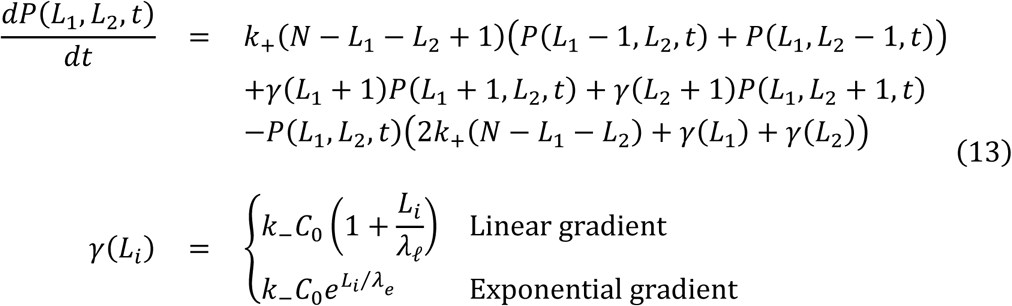

where *P*(*L*_1_, *L*_2_, *t*) is the probability that the lengths of the two filaments are *L*_1,2_ at time *t* and (*N* − *L*_1_ − *L*_2_) is the number of free monomers that are available for assembly. To obtain the steady state distribution of filament lengths, we set the left-hand side of equation (13) equal to zero and solve for the probability *P*(*L*_1_, *L*_2_), which is time independent.

From the master equation, it follows that the steady state probability distribution must satisfy the recursion relations, 

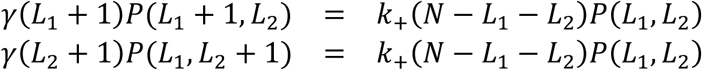

and so, the solution of the master equation in steady state is given by, 

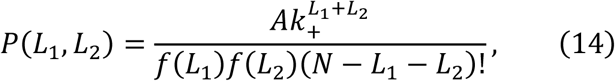

where, 

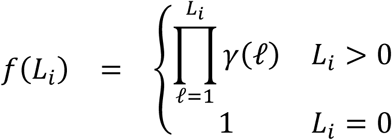

From the joint probability distribution of lengths of the two filaments, we obtain the steady state distribution of lengths of a single filament, say *L*_1_, by summing over all possible lengths of the other filament (*L*_2_). In figures 3A and 3B we plot the steady state probability distribution for a single filament, in the case of a linear and an exponential gradient of depolymerizers, using the same parameters as in figure 2. To show the effect of a finite pool on the steady state length distribution, we compare the large pool and limiting pool versions of each model in figures 3C and 3D. We see that the main effect of a finite monomer pool is that the mean lengths are smaller in this case, while the noise is almost the same in both cases. It changes from 0.007 to 0.006 for the linear gradient model, while for the exponential gradient model the difference is negligible. This conclusion does not change even when the number of monomers in the filaments is a much larger fraction of the finite pool. We explore the effect of the finite pool further in figures 3E and 3F, where we plot the steady-state lengths when a finite monomer pool is taken into account and when it is not. As the fraction of monomers in the filaments decreases the difference between the predictions of the finite and large monomer pool case also decrease, as expected.

**Figure 3:**
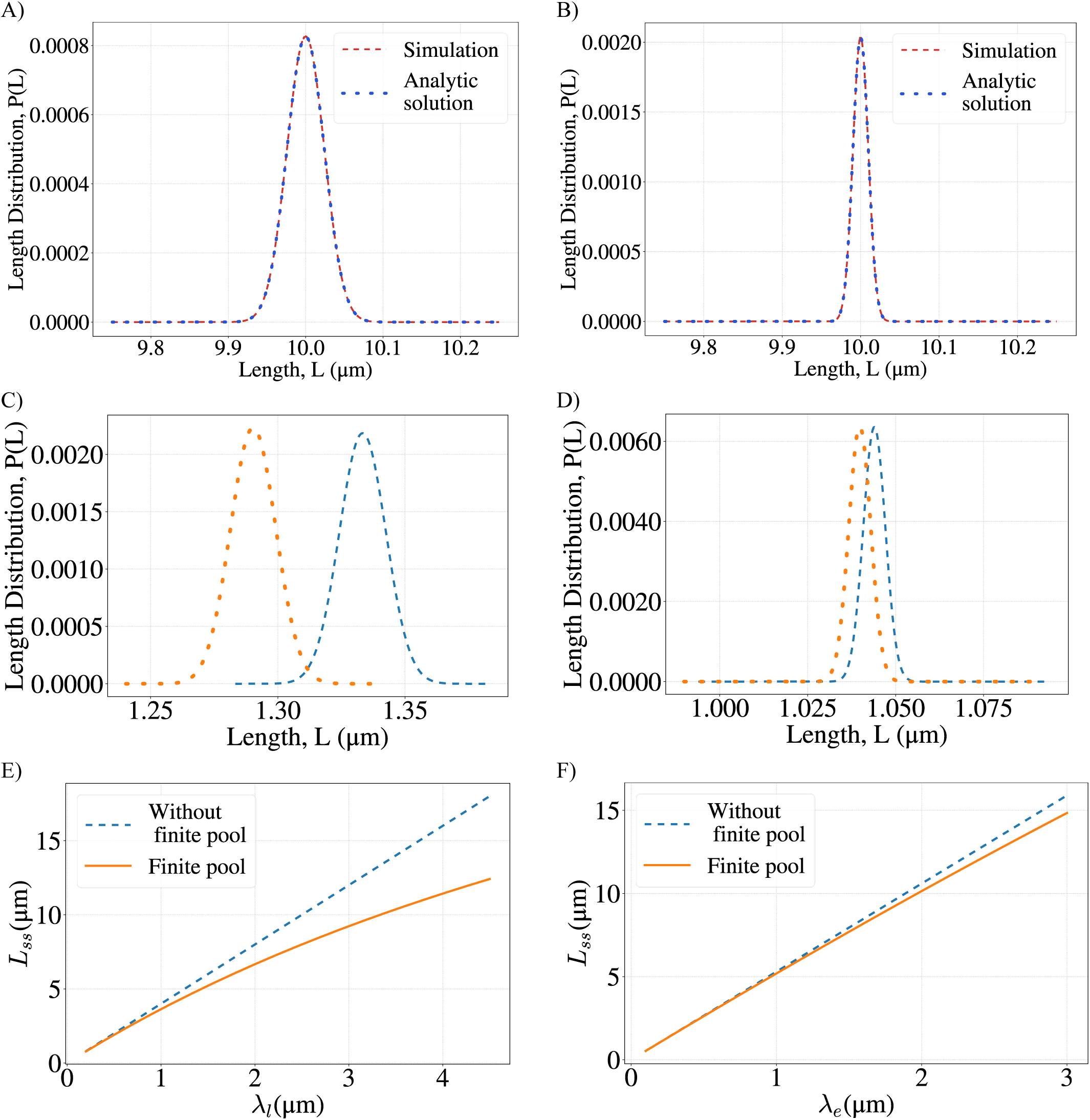
**A, B**. Marginal distributions of the filament lengths for the linear (A) and the exponential (B) gradient model with finite monomer pool. Simulation results are in agreement with the analytical solutions; see equations (15) and (17). **C, D**. Compares results for the finite pool and the large pool cases, when only a small fraction (∼2%) of monomers is in filaments. (C) corresponds to the linear, while (D) is the exponential gradient model. In both plots, the orange dashed curve represents the finite pool model and the light blue dashed curve represents the large pool model limit. As expected, the effect of the finite pool is small in this case. **E, F**. Shows the difference between steady state lengths (*L*_*ss*_) for the large monomer pool model (light blue) and the finite pool model (orange) for different values of the length scale of the gradient. Parameters: *N* = 2 × 10^6^. For **A**: *QN* = 5, *λ*_*l*_ = 6.66 × 10^4^(in monomer units); **B**: *QN* = 200, *λ*_*e*_ = 3.94 × 10^4^ (in monomer units); **C**: *QN* = 5, *λ*_*l*_ = 6.66 × 10^3^(in monomer units); **D**: *QN* = 200, *λ*_*e*_ = 3.94 × 10^3^ (in monomer units); **E**. *QN* = 5; **F**: *QN* = 200.

To further explore the consequences of the length distribution, equation (14), we examine how the mean length and the noise of the distribution depend on the two parameters of the model *QN* and *λ*, while keeping the number *N* of monomers fixed.

For the linear gradient model equation (14) can be further simplified to, 

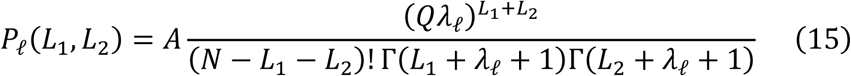

The mean and variance are not analytically tractable, but we can compute the steady state length from the system of equations *r*(*L*_SS_, *L*_SS_) = *γ*_*ℓ*_(*L*_*SS*_), which yields 

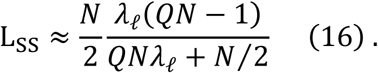

We plot this function in figure 4A (which is practically indistinguishable from the mean length).

**Figure 4:**
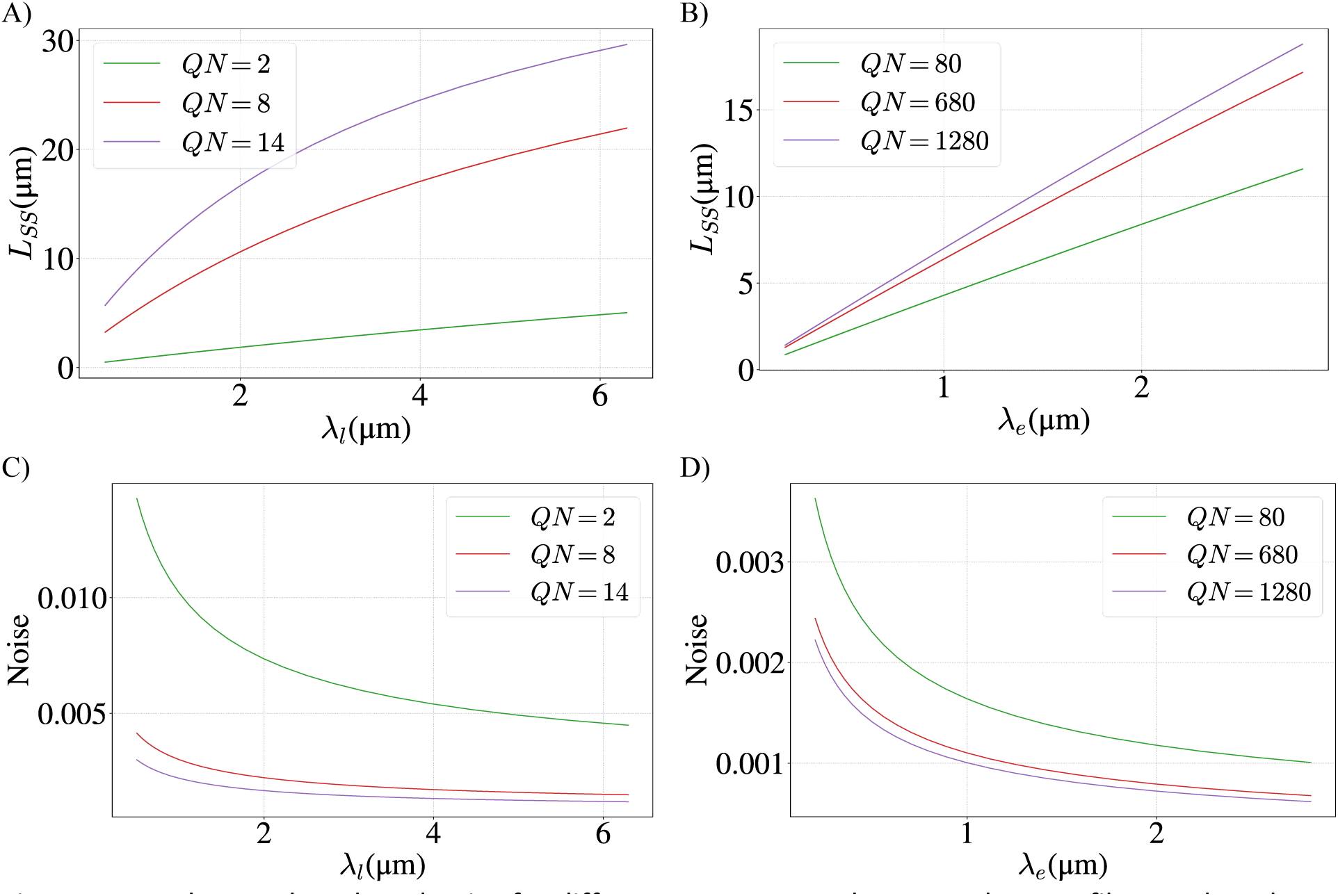
Steady state length and noise for different parameter values. Steady-state filament length for the linear (**A**) and exponential (**B**) gradient models, for different values of the dimensionless rate parameter *QN* and gradient length scale *λ*_*l,e*_. Noise of the length distribution for the linear (**C**) and exponential (**D**) gradient model for the same parameter values as in A and B. For all graphs the total number of monomers is *N* = 2 × 10^6^.

The variance cannot be computed analytically in this manner. Instead we use Equation (15) to compute the variance of the length distribution for a single filament, and in figure 4C we show how the noise of the distribution 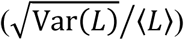 depends on the parameters *λ*_*l*_ and *Q*. We see that the noise is much less than one for a large range of parameters. This gives a measure of how well controlled the filament lengths are for different parameter values.

For the exponential gradient of depolymerizers, Equation (14) simplifies to 

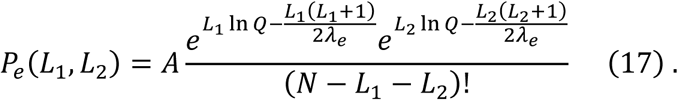

In this case we cannot obtain an analytic formula for either the steady state length or the variance. The system of equations *r*(*L*_SS_, *L*_SS_) = *γ*_*e*_(*L*_*SS*_) does not have an analytic solution but is easily solved numerically to find approximate values for the steady state length, which we find to be an excellent approximation (at most, one monomer difference) of the mean. As with the linear gradient model we find that the noise is much less than one for a range of parameters, figure 4D.

## IV. Discussion

Here we considered a simple model of length control of filaments undergoing rapid turnover of their constituent monomers, where the rate of depolymerization increases with filament length. Such length dependent depolymerization can be established in filamentous structures such as flagella and cilia, within which protein gradients are produced by a combination of directed motor transport and diffusion. If the protein being transported has a depolymerizing activity, then the accumulation of this protein at the filament tip (see figure 1) will lead to a length dependent depolymerization of the filament on which it is transported. Depending on the details of the transport, the amount of protein accumulated at the tip, and therefore its depolymerizing activity, will either depend linearly or exponentially on the filament length [20,24]. Most importantly, in the presence of multiple filaments drawing from the same pool of depolymerizing proteins, the concentration at the tip of each filament will only depend on the length of that particular filament, thereby enabling simultaneous length control of multiple filaments [18]. This is in contrast to length control by a finite monomer pool in which case only the total length of all the filaments is precisely controlled, while each individual filament’s length can vary wildly [12].

The key result of this paper are analytical expressions for the probability distributions of filament lengths when the dominant mechanism of length control is length-dependent depolymerization, due to either an exponential or a linear gradient of depolymerizing proteins. Notably, both types of protein gradients have been observed in flagella of single-cell organisms, Chlamydomonas and Giardia [19,25]. We derive analytical results for two scenarios: when the pool of filament subunits is limited (i.e., a sizable fraction of all monomers is in filamentous form in steady state), and when it is effectively infinite (i.e., the fraction of monomers in filaments is small). Estimates of model parameters based on experiments in Chlamydomonas yield steady state lengths that are in agreements with observations. Based on the same parameters our estimates of noise are two orders of magnitude smaller than the observed length fluctuations pointing to a different source of noise not accounted for by our models. Even though we only explore a two-filament system, our calculations can easily be extended to multiple filaments and the probability distributions can be found in a similar way.

A notable, qualitative difference between the linear and exponential gradient models is that in the latter case the steady state filament length is set by the length scale of the gradient and is only logarithmically dependent on the rates of polymerization and depolymerization (see Equation (6)). This makes the exponential gradient model more robust in controlling the length than the linear gradient model, for which the steady state length is more sensitive to model parameters, such as the total number of monomers and the concentration of depolymerizing proteins in the cell (see Equation (5)). Therefore, we are left with an interesting prediction of the exponential gradient mechanism, namely that orders of magnitude changes in the cytoplasmic concentration of the depolymerizing protein will lead to much smaller changes in filament length (see Figure 4B). Future experiments that combine genetic and drug manipulations of cells in order to manipulate protein gradients within the flagellum, while at the same time measuring flagellar lengths, should provide stringent experimental tests of our predictions.

## V. Acknowledgements

This work was supported by the National Science Foundation grants DMR-1610737 and MRSEC-1420382, and by the Simons Foundation. We are grateful to Lishibanya Mohapatra, Prathitha Kar, Shane McInally, Thomas Fai, and Ariel Amir for useful discussions and comments on the manuscript.

## References

1. Fletcher DA, Mullins RD. Cell mechanics and the cytoskeleton. Nature. 2010;463: 485–492. doi: 10.1038/nature08908

2. Needleman DJ, Farhadifar R. Mitosis: Taking the Measure of Spindle Length. Current Biology. 2010;20: R359–R360. doi: 10.1016/j.cub.2010.02.054

3. Mohapatra L, Goode BL, Kondev J. Antenna Mechanism of Length Control of Actin Cables. Sun SX, editor. PLoS Comput Biol. 2015;11: e1004160. doi: 10.1371/journal.pcbi.1004160

4. Marshall WF. CELLULAR LENGTH CONTROL SYSTEMS. Annu Rev Cell Dev Biol. 2004;20: 677–693. doi: 10.1146/annurev.cellbio.20.012103.094437

5. Wemmer KA, Marshall WF. Flagellar Length Control in Chlamydomonas—A Paradigm for Organelle Size Regulation. International Review of Cytology. Elsevier; 2007. pp. 175–212. doi: 10.1016/S0074-7696(06)60004-1

6. Melbinger A, Reese L, Frey E. Microtubule Length-Regulation by Molecular Motors. Phys Rev Lett. 2012;108: 258104. doi: 10.1103/PhysRevLett.108.258104

7. Johann D, Erlenkämper C, Kruse K. Length Regulation of Active Biopolymers by Molecular Motors. Phys Rev Lett. 2012;108: 258103. doi: 10.1103/PhysRevLett.108.258103

8. Good MC, Vahey MD, Skandarajah A, Fletcher DA, Heald R. Cytoplasmic Volume Modulates Spindle Size During Embryogenesis. Science. 2013;342: 856–860. doi: 10.1126/science.1243147

9. Kuan H-S, Betterton MD. Biophysics of filament length regulation by molecular motors. 2014; 28.

10. Harbage D, Kondev J. Exact Length Distribution of Filamentous Structures Assembled from a Finite Pool of Subunits. J Phys Chem B. 2016;120: 6225–6230. doi: 10.1021/acs.jpcb.6b02242

11. Chesarone-Cataldo M, Guérin C, Yu JH, Wedlich-Soldner R, Blanchoin L, Goode BL. The Myosin Passenger Protein Smy1 Controls Actin Cable Structure and Dynamics by Acting as a Formin Damper. Developmental Cell. 2011;21: 217–230. doi: 10.1016/j.devcel.2011.07.004

12. Mohapatra L, Lagny TJ, Harbage D, Jelenkovic PR, Kondev J. The Limiting-Pool Mechanism Fails to Control the Size of Multiple Organelles. Cell Systems. 2017;4: 559-567.e14. doi: 10.1016/j.cels.2017.04.011

13. Kuchka MR, Jarvik JW. Analysis of flagellar size control using a mutant of Chlamydomonas reinhardtii with a variable number of flagella. The Journal of Cell Biology. 1982;92: 170–175. doi: 10.1083/jcb.92.1.170

14. Marshall WF, Qin H, Brenni MR, Rosenbaum JL. Flagellar Length Control System: Testing a Simple Model Based on Intraflagellar Transport and Turnover. MBoC. 2005;16: 270–278. doi: 10.1091/mbc.e04-07-0586

15. Coyne B, Rosenbaum JL. FLAGELLAR ELONGATION AND SHORTENING IN CHLAMYDOMONAS. The Journal of Cell Biology. 1970;47. doi: 10.1083/jcb.41.2.600

16. Ishikawa H, Marshall WF. Ciliogenesis: building the cell’s antenna. Nat Rev Mol Cell Biol. 2011;12: 222–234. doi: 10.1038/nrm3085

17. Hendel NL, Thomson M, Marshall WF. Diffusion as a Ruler: Modeling Kinesin Diffusion as a Length Sensor for Intraflagellar Transport. Biophysical Journal. 2018;114: 663–674. doi: 10.1016/j.bpj.2017.11.3784

18. Fai TG, Mohapatra L, Kar P, Kondev J, Amir A. Length regulation of multiple flagella that self-assemble from a shared pool of components. eLife. 2019;8: e42599. doi: 10.7554/eLife.42599

19. McInally SG, Kondev J, Dawson SC. Length-dependent disassembly maintains four different flagellar lengths in Giardia. Pazour GJ, editor. eLife. 2019;8: e48694. doi: 10.7554/eLife.48694

20. Varga V, Leduc C, Bormuth V, Diez S, Howard J. Kinesin-8 Motors Act Cooperatively to Mediate Length-Dependent Microtubule Depolymerization. Cell. 2009;138: 1174–1183. doi: 10.1016/j.cell.2009.07.032

21. Niwa S, Nakajima K, Miki H, Minato Y, Wang D, Hirokawa N. KIF19A Is a Microtubule-Depolymerizing Kinesin for Ciliary Length Control. Developmental Cell. 2012;23: 1167–1175. doi: 10.1016/j.devcel.2012.10.016

22. Helenius J, Brouhard G, Kalaidzidis Y, Diez S, Howard J. The depolymerizing kinesin MCAK uses lattice diffusion to rapidly target microtubule ends. Nature. 2006;441: 115–119. doi: 10.1038/nature04736

23. Blaineau C, Tessier M, Dubessay P, Tasse L, Crobu L, Pagès M, et al. A Novel Microtubule-Depolymerizing Kinesin Involved in Length Control of a Eukaryotic Flagellum. Current Biology. 2007;17: 778–782. doi: 10.1016/j.cub.2007.03.048

24. Naoz M, Manor U, Sakaguchi H, Kachar B, Gov NS. Protein Localization by Actin Treadmilling and Molecular Motors Regulates Stereocilia Shape and Treadmilling Rate. Biophysical Journal. 2008;95: 5706–5718. doi: 10.1529/biophysj.108.143453

25. Chien A, Shih SM, Bower R, Tritschler D, Porter ME, Yildiz A. Dynamics of the IFT machinery at the ciliary tip.: 25.

26. Gorovsky MA, Carlson K, Rosenbaum JL. Simple method for quantitive densitometry of polyacrylamide gels using fast green. Analytical Biochemistry. 1970;35: 359–370. doi: 10.1016/0003-2697(70)90196-X

27. Johnson KA, Rosenbaum JL. Polarity of Flagellar Assembly in Chlamydomonas. The Journal of Cell Biology. 1992;119: 1605–1611.

28. Song L, Dentler WL. Flagellar Protein Dynamics in *Chlamydomonas*. J Biol Chem. 2001;276: 29754–29763. doi: 10.1074/jbc.M103184200

29. Marshall WF, Rosenbaum JL. Intraflagellar transport balances continuous turnover of outer doublet microtubules. The Journal of Cell Biology. 2001;155: 405–414. doi: 10.1083/jcb.200106141

30. Bayley PM, Sharma KK, Martin SR. Microtubule dynamics in vitro. Mod cell biol. 1994;13: 111–137.

31. Craft JM, Harris JA, Hyman S, Kner P, Lechtreck KF. Tubulin transport by IFT is upregulated during ciliary growth by a cilium-autonomous mechanism. Journal of Cell Biology. 2015;208: 223–237. doi: 10.1083/jcb.201409036

32. Piperno G, Luck DJL. Microtubular proteins of Chlamydomonas reinhardtii. An immunochemical study based on the use of an antibody specific for the beta-tubulin subunit. Journal of Biological Chemistry. 1977;252: 383–391.

